# Compound heterozygosity of the gamma-glutamyl carboxylase mutants V255M and S300F causes pseudoxanthoma elasticum-like disease through impaired processivity

**DOI:** 10.1101/668574

**Authors:** Mark A. Rishavy, Kevin W. Hallgren, Kurt W. Runge, Kathleen L. Berkner

## Abstract

Vitamin K-dependent (VKD) protein activities require carboxylated Glus (Glas) generated by the gamma-glutamyl carboxylase. Some carboxylase mutations cause severe bleeding, while others cause pseudoxanthoma elasticum (PXE)-like associated with excessive calcification. How carboxylase mutations cause PXE-like was unknown. We analyzed two mutants (V255M and S300F) whose compound heterozygosity causes PXE-like. Substrates derived from VKD proteins important to calcification (MGP) or clotting (factor IX) were studied, which contained the Gla domain and exosite-binding domain that mediates carboxylase binding. Surprisingly, the V255M mutant was more active (4-5 fold) than wild type carboxylase, while S300F activity was low. The V255M results suggested faster substrate release, which could impact carboxylase processivity, where the carboxylase remains bound to VKD proteins throughout multiple Glu to Gla conversions. To assess mutant processivity, we performed a novel challenge assay in which MGP-carboxylase and factor IX-carboxylase complexes were reacted in the presence of excess challenge VKD peptide. Tight complexes between VKD proteins and wild type carboxylase excluded access of the challenge peptide during the carboxylation of VKD protein in the complex. In contrast, VKD protein complexes with V255M or S300F allowed promiscuous access of challenge peptide. Both mutants therefore impair processivity. Most of the V255M product was carboxylated challenge peptide, which could explain mild PXE-like observed in the proband’s mother and aunt. Both have wild type and V255M carboxylase alleles; however, higher V255M production of a potentially defective MGP product could account for their phenotype. The results are an important advance in understanding why carboxylase mutations cause the PXE-like disease.

## INTRODUCTION

Pseudoxanthoma elasticum (PXE) is an autosomal recessive disorder associated with ectopic mineralization in soft connective tissues, notably skin, eyes and arteries (1). Classic PXE is caused by mutations in ABCC6, a transmembrane efflux pump that transports unknown metabolite(s) into the circulation (1). PXE-like disease is also observed in patients with mutations in the gamma-glutamyl carboxylase (2). A single carboxylase modifies ∼16 vitamin K-dependent (VKD) proteins, which require the modification for activities that include hemostasis, regulation of calcification, apoptosis, growth control and signal transduction (3). The VKD protein matrix Gla protein (MGP) is a major regulator of calcification (4), and defective modification by the carboxylase is observed in both the PXE and PXE-like diseases (5). Approximately half of the VKD proteins are important to hemostasis, and PXE-like is distinct from PXE in being associated with a mild defect in multiple vitamin K-dependent (VKD) coagulation factor activities. Mild bleeding contrasts what observed in a second disease caused by carboxylase mutants, i.e. vitamin K clotting factor deficiency (VKCFD1), where mutations that substantially decrease carboxylase activity cause a hemorrhagic response that can be life threatening (6).

Why carboxylase mutations cause the PXE-like disease is unknown, and indicates the importance of understanding how the carboxylase modifies VKD proteins. The carboxylase reaction is complex and involves several substrates: vitamin K and oxygen are used to add CO_2_ to Glu to generate carboxylated Glu (Gla)(Fig. 1A). Reduced vitamin K is required, and is provided by a second enzyme, the vitamin K oxidoreductase (VKORC1)(7,8). Clusters of Glus are carboxylated in the Gla domain of VKD proteins, which results in the generation of a calcium-binding module that is important for activity. The Glus are targeted to the carboxylase by an exosite-binding domain that mediates high affinity binding of the VKD proteins to the carboxylase, and which also activates Glu catalysis (Fig. 1B). In most cases, the exosite binding domain is a propeptide (as in Fig. 1B) that is cleaved subsequent to carboxylation. The carboxylation of multiple Glus occurs through a processive mechanism (9,10). We found that VKD substrate binding to the carboxylase resulted in the complete modification of the VKD protein even in the presence of a pool of VKD proteins with the potential to compete for carboxylase binding (10). Carboxylase processivity is significant because tissues express multiple VKD proteins. Liver, for example, expresses seven hemostatic VKD factors as well as nonhemostatic VKD proteins (11), and the affinities of the exosite-binding domains in these proteins vary widely (12). These proteins compete for one carboxylase, and a processive mechanism ensures full modification.

**Figure 1.**
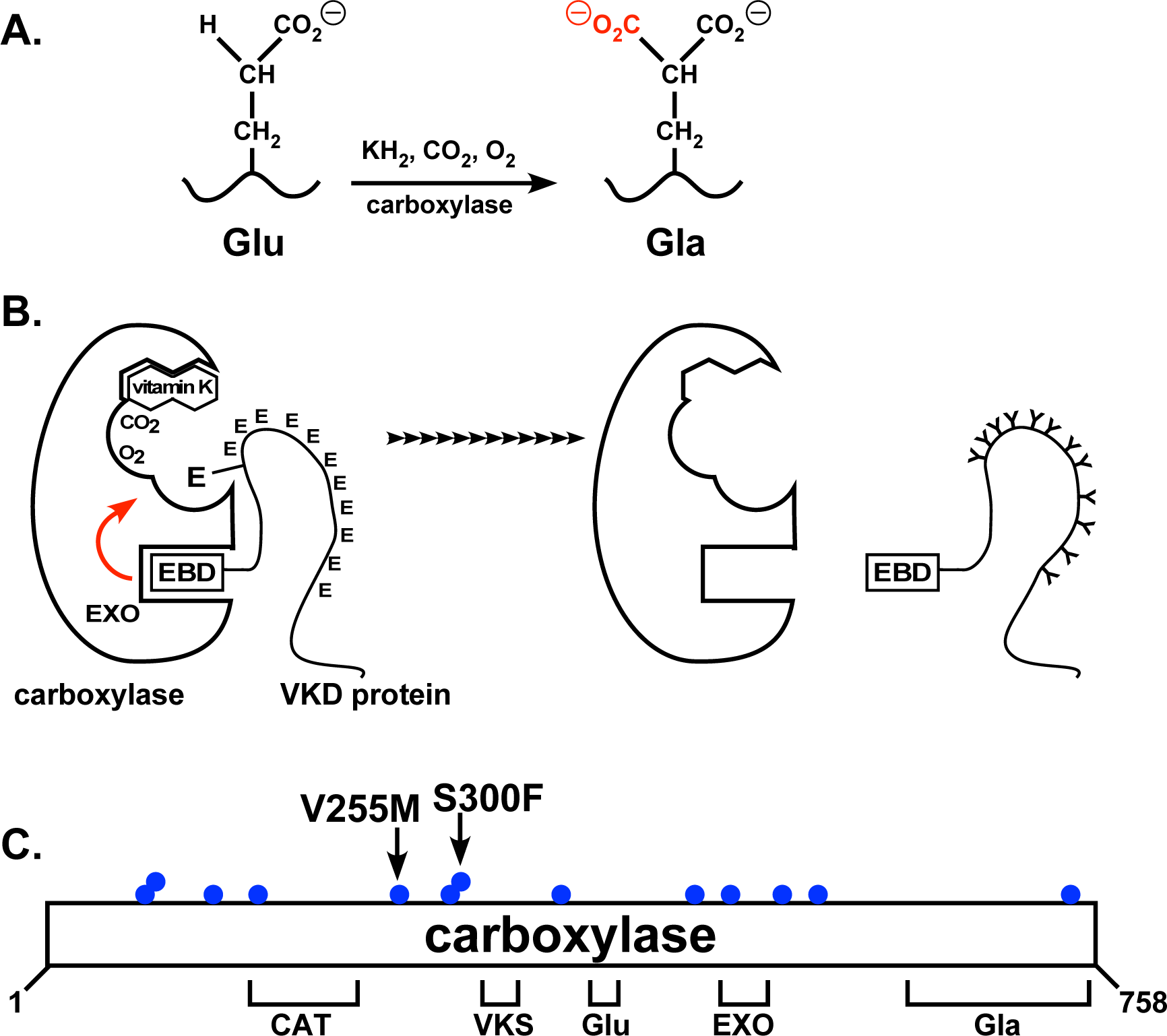
Mutations in the carboxylase cause the PXE-like disease. **A.** The carboxylase converts Glu to carboxylated Glu (Gla), using the oxygenation of reduced vitamin K (KH_2_) to drive CO_2_ addition to Glu. **B.** VKD proteins are targeted to the carboxylase through an exosite-binding domain (EBD) that mediates high affinity binding through the carboxylase exosite (EXO), and also activates Glu catalysis (red arrow). Multiple Glu residues are converted to Gla (Y) by a processive mechanism in which a single binding event between the carboxylase and VKD protein results in full carboxylation (10). **C.** Known functional regions of the carboxylase include those facilitating catalysis (CAT), VKD protein binding (VKS), Glu binding (Glu), and the exosite (EXO) and carboxylase Gla domain (GLA). Most of the residues whose mutations cause PXE-like reside in regions where the function of the carboxylase is unknown. This study shows that the V255M and S300F mutants impair carboxylase processivity.

Only some of the functional regions in the carboxylase have been identified. The known residues include ones important to catalysis, Glu-binding and VKD protein binding (Fig. 1C)(13). Binding sites for other substrates, e.g. vitamin K, are unknown, as are residues that regulate the reaction. The functional roles of most of the residues whose mutations cause the PXE-like disease are unknown, making it difficult to understand how the mutations cause the PXE-like disorder. A mutation in Asp153 underscores this point. A D153G mutation in a PXE-like patient has a larger impact on the carboxylation of MGP than a hemostatic VKD protein (14), but the reason for the differential effect is unknown. The result would suggest that Asp153 is part of the exosite that recognizes VKD proteins, however Asp153 resides instead within the catalytic domain (Fig. 1A).

We pursued a study on a patient with compound heterozygosity of the carboxylase mutations V255M and S300F (15). The patient had severe skin manifestations that included mineralization of elastin fibers, as well as defective clotting and poor carboxylation of MGP in skin. A test for activity in an assay that measures the carboxylation of a small Glu-containing substrate could not explain why the mutations caused the disease. Novel assays were therefore developed in the current study to more fully assess the complex carboxylase reaction. The assays included analyzing large peptides derived from VKD proteins important to hemostasis (factor IX (fIX)) or calcification (MGP). These substrates contained the Gla domain and exosite-binding domain, and were used to study carboxylase turnover (i.e. the binding, catalysis and release that occurs during one cycle of VKD protein carboxylation). Other studies used complexes between the carboxylase and full-length fIX or MGP to analyze carboxylase processivity. Surprisingly, V255M produced more carboxylated VKD product than wild type carboxylase, which we showed is due to impaired processivity. In the case of wild type carboxylase, a tight complex with VKD protein excluded the access of external VKD proteins while the VKD protein in the complex underwent carboxylation. In contrast, the active sites of both the V255M and S300F mutants were promiscuous, allowing inappropriate access of VKD protein. V255M produced most of the carboxylated product resulting from this unregulated access of VKD protein. Dysregulation of processivity may generate undercarboxylated MGP that is defective, explaining why the mutations cause the disease. The studies reveal the first functional understanding for the underlying mechanism of the PXE-like disorder.

## RESULTS

### The V255M mutant shows impaired exosite activation

The carboxylase V255M and S300F mutants whose compound heterozygosity caused PXE-like (15) were individually analyzed after expression in insect cells that lack endogenous carboxylase components. VKD protein binding to the carboxylase exosite activates catalysis (Fig. 1B). This activation was studied using two peptides derived from either the Gla domain or exosite-binding domain (EBD)(Fig. 2A). Purified carboxylase variants (Fig. 2B) were incubated with a Glu-containing peptide from the Gla domain (FLEEL) in the presence or absence of EBD peptide. In most VKD proteins, the EBD peptide is a propeptide that is removed subsequent to carboxylation, and the factor X EBD was used in the assay. Carboxylation was monitored by quantitating incorporation of ^14^C-CO_2_ into the FLEEL peptide. In the absence of the factor X EBD peptide, the wild type carboxylase showed significant activity, while the activity of the V255M carboxylase was very low and that of the S300F carboxylase was undetectable (Fig. 2C). In the presence of the factor X EBD peptide, the S300F carboxylase had barely detectable activity while the V255M carboxylase showed activity similar to that of wild type carboxylase (Fig. 2C). Activation of the V255M carboxylase (∼80-fold) was therefore much higher than that of wild type carboxylase (∼4-fold). Similar results were obtained with an EBD peptide derived from MGP (data not shown).

**Figure 2.**
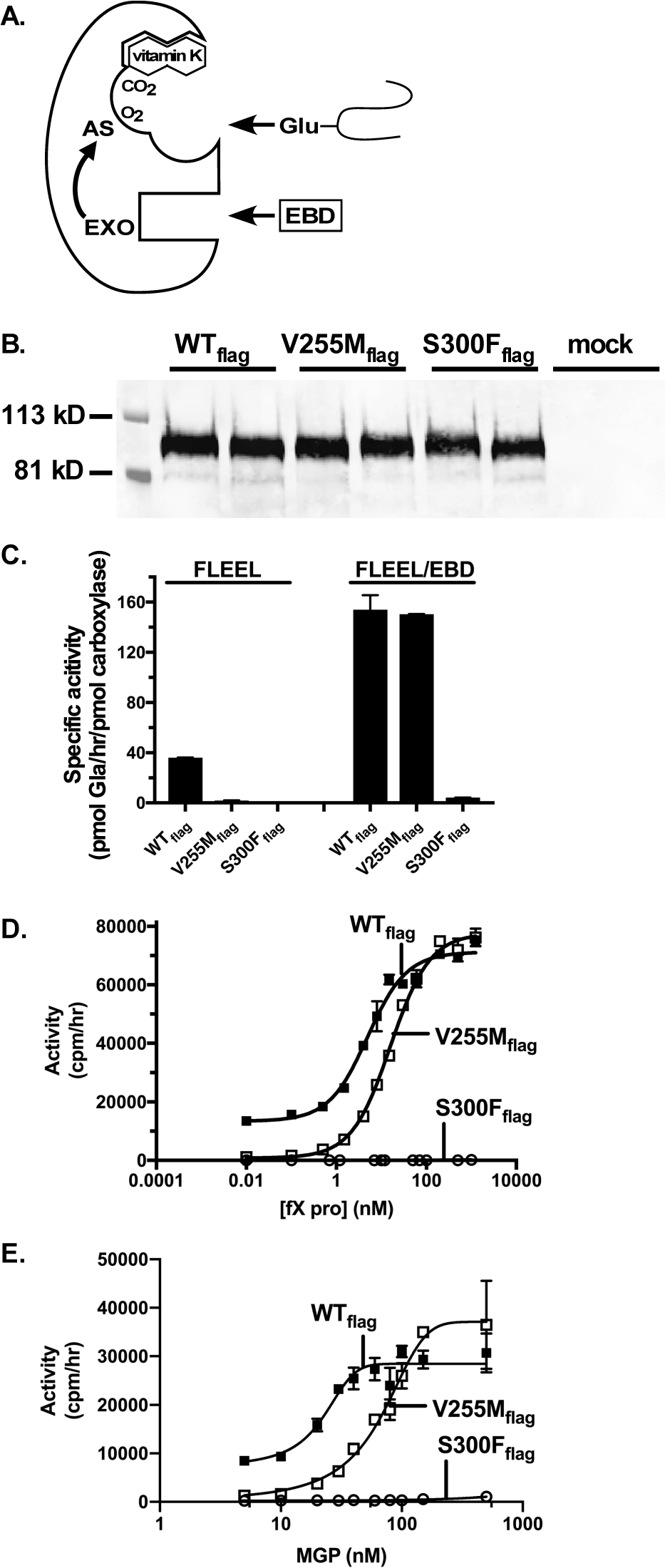
An assay for exosite activation of catalysis reveals that V255M carboxylase has impaired interaction with the exosite binding domain and S300F carboxylase has poor catalysis. **A.** VKD proteins contain two domains important to carboxylation: the Gla domain containing Glu residues that are converted to carboxylated Glu (Gla) residues, and the exosite binding domain (EBD) that mediates high affinity binding to the carboxylase exosite (EXO) and activation of catalysis in the active site (AS). Carboxylation and activation of catalysis were studied using peptides derived from each domain. **B.** Wild type (WT) and mutant carboxylase variants were expressed in insects cells, purified and then analyzed by Western to quantitate enzyme levels. **C.** Carboxylase variants were incubated with a Glu-containing peptide (FLEEL) in the presence or absence of the EBD from factor X, and ^14^C-CO_2_ incorporation into FLEEL was determined and normalized to the amount of carboxylase. **D-E.** Exosite activation of Glu catalysis for each carboxylase variant was studied with varying amounts of EBD peptides derived from factor X (D) or MGP (E). The V255M carboxylase required higher concentrations of EBD for full activation, indicating impaired interaction, while the S300F carboxylase showed almost no activity in these assays.

The activation assay (Fig. 2C) used saturating concentrations of EBD peptide (10 µM), and further analysis was performed using lower concentrations. The different carboxylase enzymes were incubated with FLEEL and varying amounts of either the factor X EBD peptide or the MGP EBD peptide, and the amount of ^14^C-CO_2_ incorporated into FLEEL was monitored. Interestingly, the V255M carboxylase required higher concentrations of either EBD peptide for activation compared to wild type carboxylase (Figs. 2D, E). For example, with the MGP EBD peptide, half-maximal activation was observed at 80 nM for V255M versus 20 nM for wild type carboxylase (Fig. 2E). These dose-response experiments also confirmed that the EBD peptide caused a much larger fold activation with V255M than wild type carboxylase. In contrast, the S300F carboxylase only showed trace levels of carboxylation in the presence of the EBD peptide (Figs. 2D, E). These results suggested that the V255M carboxylase was impaired in the interaction with EBDs of VKD proteins, which was pursued by analyzing different types of VKD substrates.

### The V255M carboxylase shows surprisingly higher VKD protein carboxylation than wild type enzyme

The carboxylase variants were analyzed using synthetic peptides that mimic VKD proteins, i.e. with the Gla domain covalently linked to the EBD. The EBD mediates high affinity binding that lowers the K_m_ for Glu residues from mM to µM levels (10,16,17). Covalent linkage of the EBD to the Gla domain tethers VKD proteins to the carboxylase to allow multiple Glu modifications (10). Therefore, carboxylation of substrates where the Gla domain is covalently linked to the EBD involves three steps to turnover the carboxylated product: binding, catalysis and release (shown for MGP in Fig. 3A). Previous work has shown that release is much slower than the other two steps (9,10,18). Therefore, the results from assaying VKD substrates in which the Gla domain and EBD are linked is very different from when these domains are separated. With a peptide where both domains are linked, the carboxylated substrate must be released before a new substrate binds to undergo carboxylation (Fig. 3A). With separate peptides, the EBD peptide can remain bound while the low affinity Glu substrate moves in an out of the active site (Fig. 2A).

**Figure 3.**
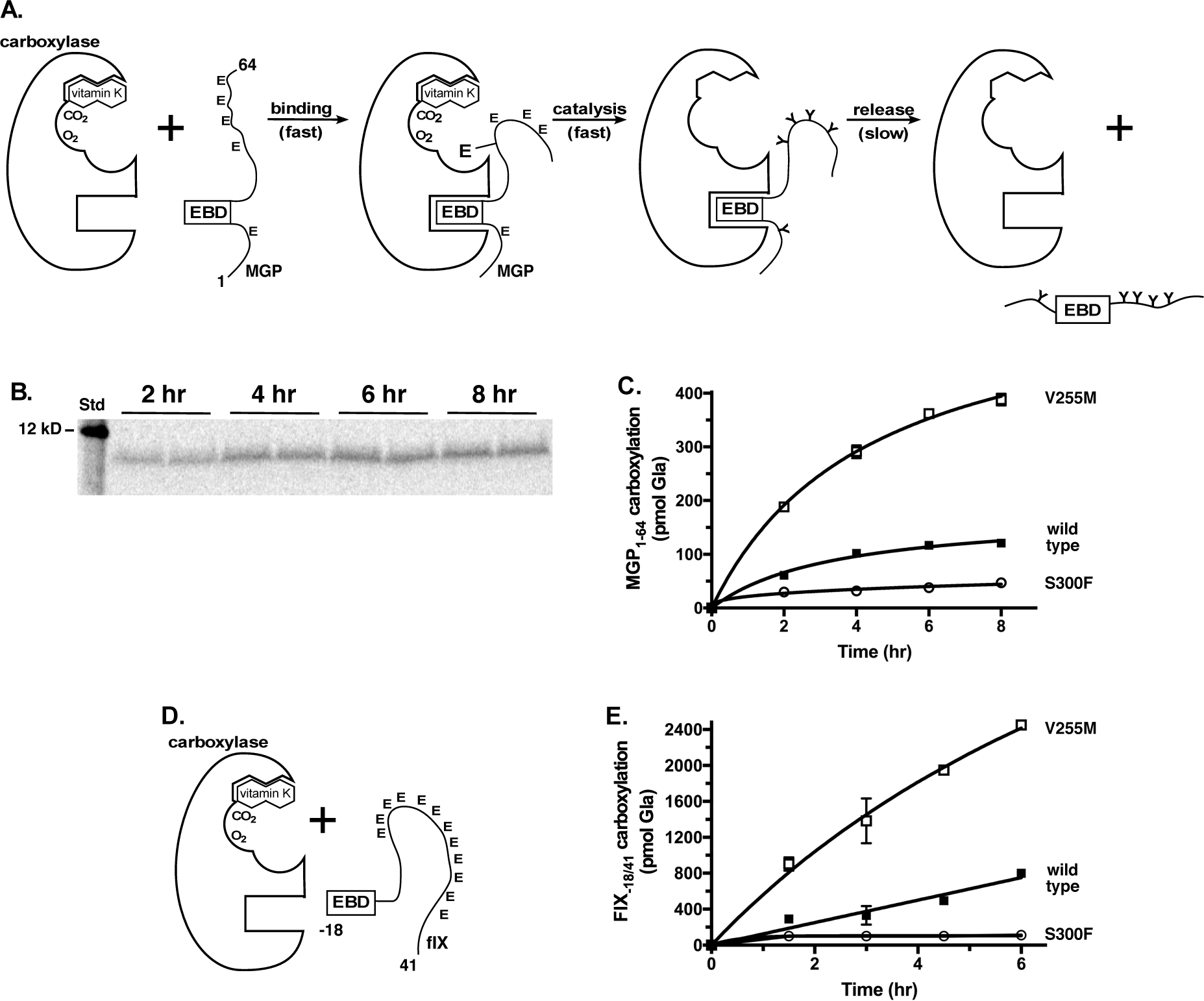
The V255M variant shows higher levels of carboxylation than the wild type carboxylase. **A.** Carboxylation was studied using a peptide (MGP_1-64_) consisting of the first 64 residues of MGP. The peptide contains all 5 Glu residues (E) that are normally carboxylated to Gla (Y) and the exosite binding domain (EBD) that mediates carboxylase binding. Carboxylation of the peptide occurs in 3 steps: binding, catalysis and release. Release is much slower than the other two steps. **B.** Carboxylation was assayed by monitoring ^14^C-CO_2_ incorporation into MGP_1-64_. Products from duplicate reactions were subjected to gel electrophoresis, followed by PhosphorImager analysis to quantitate the amount of Gla produced, using a ^14^C standard (Std) run on the same gel. **C.** The V255M carboxylase variant showed higher levels of MGP_1-64_ carboxylation than wild type carboxylase, while the activity of the S300F variant was low. **D.** Carboxylation was also studied using a peptide derived from factor IX (fIX_-18/41_), which has an 18 amino acid EBD that is cleaved from mature fIX after carboxylation, and a Gla domain containing 12 Glu (E) residues. **E.** The V255M variant showed higher levels of fIX_-18/41_ carboxylation than wild type carboxylase, while the S300F variant showed much lower levels.

Carboxylase variants were analyzed using truncated MGP (MGP_1-64_) that contains the first 64 of the 93 residues present in MGP. This peptide contains all 5 Glus that undergo carboxylation covalently linked to the propeptide that is cleaved after carboxylation. MGP is distinct from other VKD proteins in that the EBD is not a propeptide that is later removed, but instead is present in the mature protein (Fig. 3A)(13). MGP had not been tested as a substrate for carboxylation in biochemical assays, and analysis of the MGP_1-64_ peptide revealed a K_M_ (0.5 µM, data not shown) similar to what has been observed with other substrates where the EBD and Glu residues are covalently linked (10,16,17). To assay mutant carboxylases, the carboxylase variants were incubated in a reaction containing ^14^C-CO_2_, vitamin K hydroquinone (KH_2_) and saturating concentrations of MGP_1-64_ peptide (5 µM). Reaction products were analyzed by gel electrophoresis alongside a ^14^C standard, and the amount of MGP_1-64_ carboxylation was quantitated by PhosphorImager analysis (shown for wild type carboxylase in Fig. 3B, C). The rate of MGP carboxylation was compared to the amount of carboxylase in the reaction, which was quantitated both by Western analysis and by an epoxidase assay. This comparison allowed the calculation of the turnover number for MGP_1-64_, i.e. the time for one round of binding, catalysis and release. Turnover was one MGP_1-64_ per 20 min, similar to that previously determined for factor IX (10). This turnover was ∼60-fold slower than the FLEEL catalytic rate (∼3 peptides/min, Fig. 2C), consistent with previous studies showing that release is much slower than catalysis (9,10).

Interestingly, the V255M carboxylase produced much higher levels of carboxylated MGP_1-64_ than the wild type enzyme (Fig. 3C). At the end of the reaction, the amount of product produced by the V255M carboxylase was 4-fold higher than the equivalent amount of wild type enzyme. A second substrate derived from the clotting protein factor IX (fIX) was also analyzed. The EBD in fIX is an 18 amino acid propeptide that is cleaved off after carboxylation (19). The substrate, fIX_-18/41_, contained the EBD covalently linked to the Gla domain, which contains 12 Glus that undergo carboxylation (Fig. 3D). Increased carboxylation of the fIX_-18/41_ peptide by the V255M carboxylase compared to wild type enzyme was also observed (Fig. 3E). These activities on two structurally diverse VKD protein substrates showed that increased carboxylation was a general property by the V255M enzyme.

The rate of carboxylation was probed further using an assay in which catalysis can be determined independent of the binding and release steps. Catalysis was assayed by monitoring the carboxylation of VKD protein-carboxylase complexes (Fig. 4A). To generate these complexes, full-length MGP and fIX were each coexpressed with individual carboxylase variants in insect cells (Fig. 4B, C). The complexes were then separated from free carboxylase by immunoadsorption to immobilized antibody. MGP_flag_-carboxylase complexes were isolated with anti-FLAG antibody, and fIX-carboxylase complexes were isolated with anti-fIX heavy chain antibody (which binds distant from the Gla domain). We have previously shown that this immobilization does not interfere with the carboxylase reaction (10). The complexes were incubated with ^14^C-CO_2_ and KH_2_, and reaction products were then analyzed by gel electrophoresis and PhosphorImager. The V255M and wild type carboxylases showed similar rates of catalysis with both fIX and MGP (Fig. 4D, E). Therefore, the increased rate of carboxylation for V255M (Fig. 3C) was not due to altered catalysis, strongly suggesting that V255M shows increased carboxylation due to faster release.

**Figure 4.**
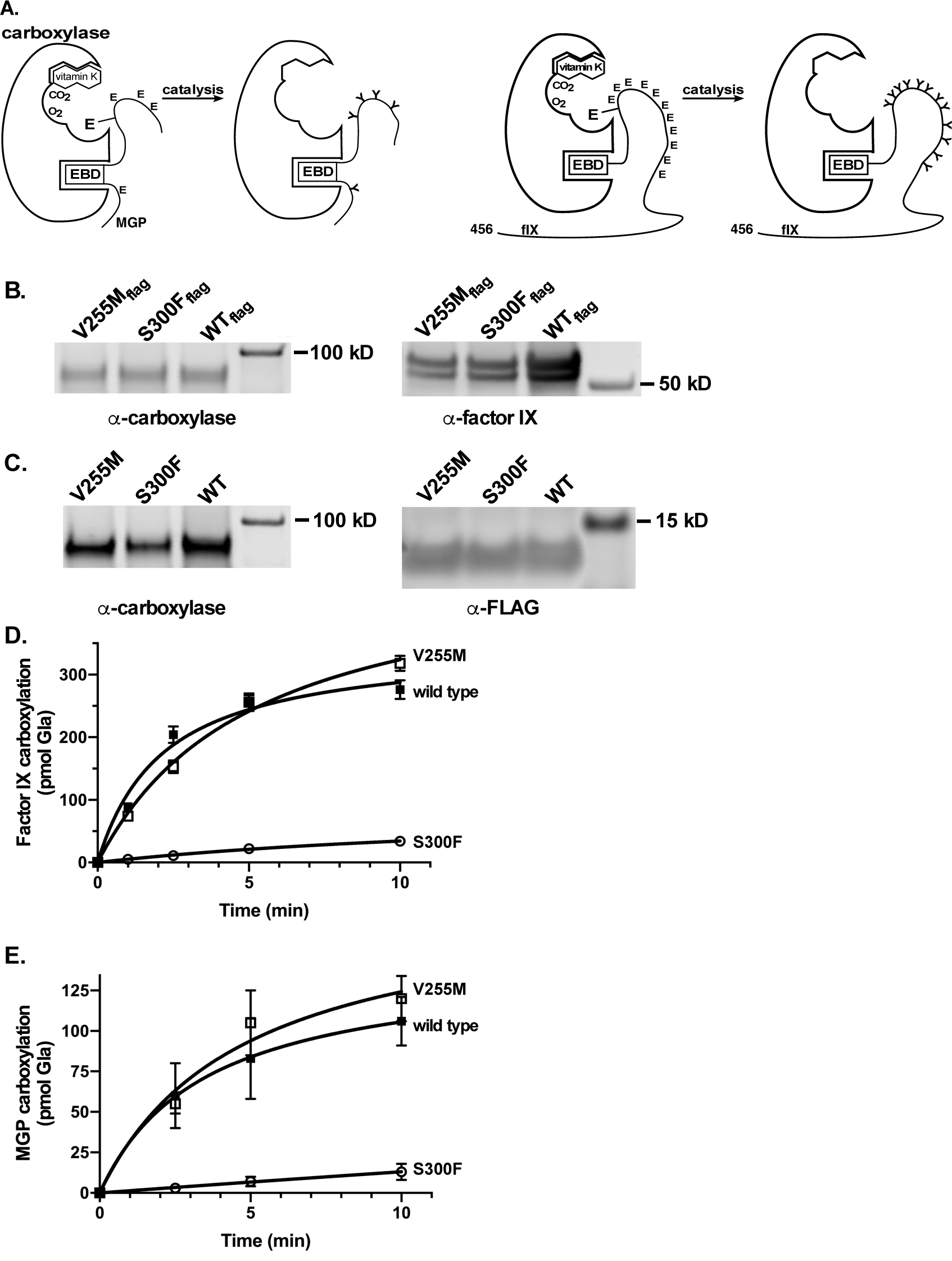
The V255M and wild type carboxylases have similar rates of VKD protein carboxylation, while the rate of S300F is much lower. **A.** MGP_flag_-carboxylase and fIX-carboxylase complexes were generated and then tested in a reaction to determine the rate of catalysis. **B-C.** Complexes were generated by co-expressing full length fIX or MGP_flag_ with the carboxylase variants in insect cells. The levels of carboxylase variant and fIX (B) or carboxylase variant and MGP_flag_ (C) were analyzed by western analysis. The fIX-carboxylase complexes were isolated from free carboxylase using an antibody against the fIX heavy chain that is distal to Gla domain, and the MGP_flag_-carboxylase complexes were isolated using anti-FLAG antibody. **D-E.** The VKD-carboxylase complexes were incubated with ^14^C-CO_2_ and vitamin K hydroquinone (KH_2_) for varying times and the reaction products were analyzed by gel electrophoresis and PhosphorImager to monitor ^14^C-CO_2_ into fIX (D) or MGP (E), using a ^14^C standard run in parallel to quantitate Gla production.

In contrast to V255M, the S300F carboxylase showed a substantially lower rate of catalysis. The S300F carboxylase, which showed little or no FLEEL carboxylation (Fig. 2C), showed clearly detectable MGP_1-64_ and fIX_-18/41_ carboxylation (Fig. 3D, E). One possible explanation for these differences was that poor catalysis of MGP_1-64_ and fIX_-18/41_ was masked because release is rate limiting for substrates with a covalently linked EBD. Analysis of the S300F carboxylase complexes with MGP or fIX revealed substantially slower rates of catalysis (Fig. 4D, E), indicating that the S300F mutation impairs carboxylase activity in a very different manner than the V255M mutation.

### Processive carboxylation of factor IX is impaired in the V255M mutant

The results with the fIX_-18/41_ and MGP_1-64_ substrates suggested faster V255M release of the carboxylated product, raising the possibility that carboxylase processivity was disrupted. Processive carboxylation ensures that the VKD proteins remain bound to the carboxylase for a sufficiently long time to allow full carboxylation of the multiple Glu residues in the Gla domain. However, a mutant enzyme that shows faster release may impair processivity by releasing a product that is not fully modified. V255M and S300F carboxylases were therefore tested for processivity, using an assay that we had previously developed (10). A preformed VKD protein-carboxylase complex is incubated in the presence of a challenge VKD peptide that is present in large excess over the complex (Fig. 5A), and carboxylation of both forms is monitored. The challenge peptide will compete with VKD protein that is prematurely released from the complex, resulting in decreased carboxylation of this VKD protein. We previously showed that premature release did not occur in a fIX-carboxylase complex containing wild type carboxylase, as fIX carboxylation was the same in the presence or absence of challenge VKD protein (10). This approach was extended to studying the V255M and S300F carboxylase variants, and both fIX and MGP carboxylation were analyzed.

**Figure 5.**
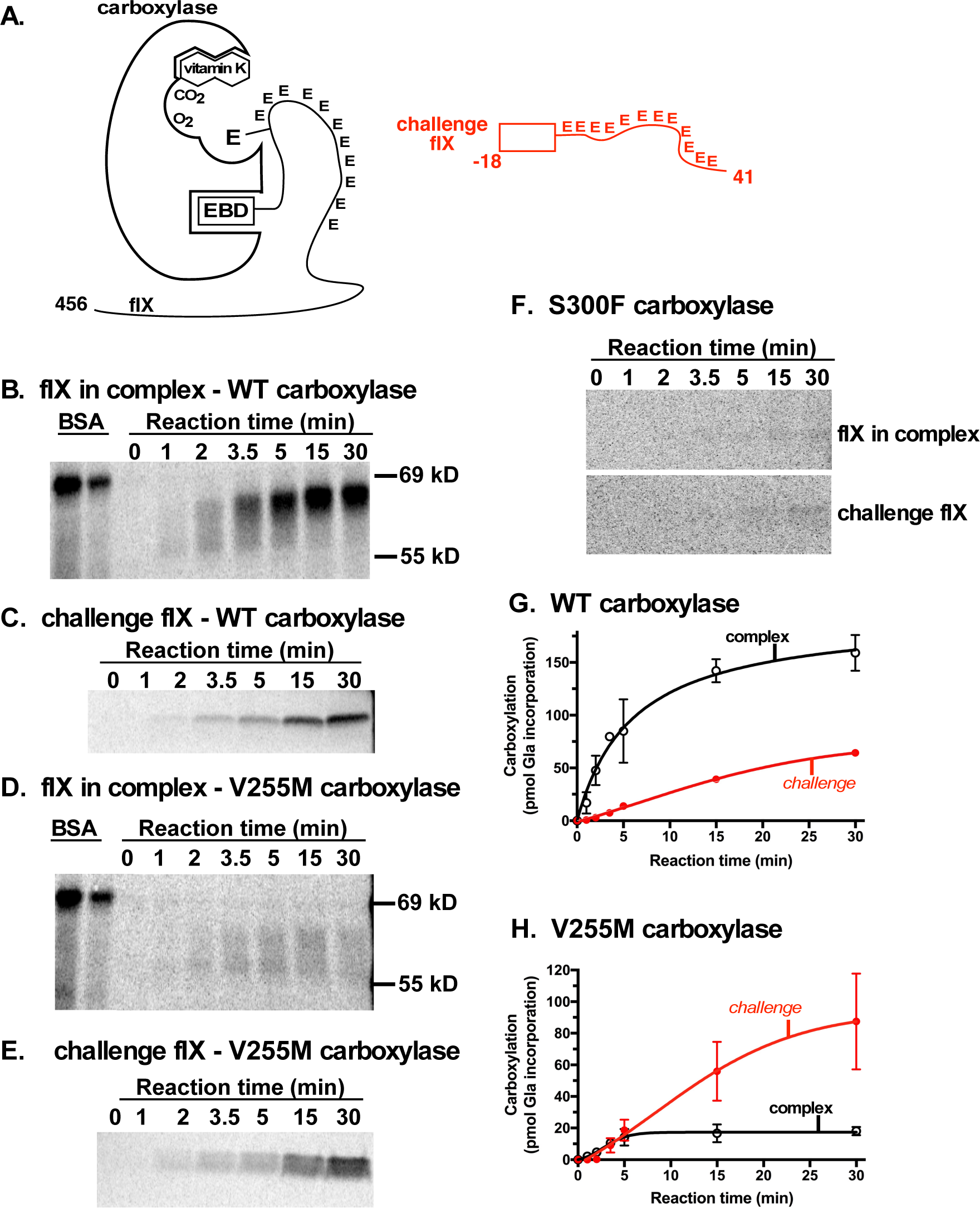
Processive carboxylation of factor IX is impaired in the V255M mutant. **A.** A reaction containing a fIX-carboxylase complex and a 50-fold excess of the fIX_-18/41_ challenge peptide was performed, and the carboxylation of both full length fIX and fIX_-18/41_ were monitored. The fIX-carboxylase complex was generated and purified as described in the legend to Figure 4. The fIX_-18/41_ challenge peptide is the same peptide analyzed for carboxylation in Fig. 3E. **B-F.** The complexes between fIX and wild type (WT) carboxylase (B, C) or V255M (D, E) or S300F (F) were incubated in reactions containing ^14^C-CO_2_, KH_2_ and fIX_-18/41_, and the products were analyzed by gel electrophoresis and PhosphorImager. Two different gels were used to monitor fIX and fIX_-18/41_ carboxylation due to the large differences in size between the two substrates. ^14^C-BSA and ^14^C-peptide standards were analyzed on the same gel to quantitate carboxylation products. **G-H.** Duplicate experiments for wild type (G) and V255M (H) carboxylases were plotted, revealing a much higher level of carboxylated fIX_-18/41_ challenge peptide with V255M than wild type carboxylase. The S300F activity was too low to quantitate.

The challenge assay for fIX-carboxylase complexes used fIX_-18/41_ as the challenge peptide. This peptide was found to be a suitable challenge substrate in potentially competing with fIX in the complex because its kinetic values (K_M_, k_cat_) were the same as propeptide-containing full-length fIX (data not shown). The fIX-carboxylase complex was generated by coinfection of insect cells with baculoviruses expressing fIX and individual carboxylase variants, as in Fig. 3. The cells were cultured in the absence of vitamin K, and so fIX in the complex was uncarboxylated. The cells contained some free carboxylase (10-20%), which would complicate the analysis, and the complexes were therefore separated from free carboxylase by adsorption to anti-fIX antibody resin. The antibody was directed against the heavy chain of fIX that is distant from the Gla domain and does not react with the fIX_-18/41_ challenge peptide (data not shown). We previously showed that immobilization did not interfere with carboxylation, as the rate of fIX-carboxylase carboxylation was only slightly slower than that of the free complex (10).

Carboxylation of the fIX-wild type carboxylase complex was assayed in a reaction mixture containing ^14^C-CO_2_, KH_2_ and fIX_-18/41_ that was present in a 50-fold excess over complex. Carboxylation was monitored over time by analyzing the reaction products by gel electrophoresis and PhosphorImager (Fig. 5B, C). The image for fIX in the complex was informative, as fIX underwent a mobility shift upon carboxylation of the 12 Glus. By the first timepoint, a heterogenous mixture of fIX forms was observed (Fig. 5B). Over time, a discrete form was observed and carboxylation reached a plateau (Fig. 5G). This plateau represented full carboxylation, as determined by comparing the amount of ^14^C-CO_2_-incorporation into fIX to the concentration of carboxylase in the complex, which was determined by an epoxidase assay. Therefore, the challenge peptide did not disrupt carboxylation of the fIX in the complex. Carboxylation of the fIX_-18/41_ challenge peptide occurred largely after the carboxylation of fIX in the complex (Fig. 5G). The results are similar to what was observed in the previous studies that established carboxylase processivity (10). While the previous work used saturating amounts of KH_2_ (150 µM)(10), the current study used lower KH_2_ concentrations (10 µM), which did not affect processivity (Fig. 5B, G).

Some carboxylated fIX_-18/41_ was observed before carboxylation of fIX in the complex was completed (Fig. 5B, C, G). This result is likely due to the nonsynchronous carboxylation of individual fIX-carboxylase complexes. Thus, carboxylation with the Gla domain requires intramolecular movement to reposition Glu residues for catalysis, which may not be identical between individual complexes. The fIX-carboxylase complexes will therefore complete fIX carboxylation at different times, and those complexes which complete modification earliest would begin carboxylating the challenge peptide before the remaining complexes have finished.

The results with the fIX-V255M carboxylase complex were strikingly different. The extent of fIX carboxylation was ∼10 lower in the fIX-V255M complex (Fig. 5D, H) compared to the fIX-wild type complex (Fig. 5 B, G), even though both experiments used equivalent amounts of complexes. The fIX-V255M complex produced a heterogeneous mixture of carboxylated fIX forms (Fig. 5D) that did not transition into the discrete form observed with wild type carboxylase (Fig. 5B). However, the V255M carboxylase complex generated much more carboxylated fIX_-18/41_ challenge peptide than the wild type carboxylase (compare Figs. 5E, H vs. Figs. 5C, G). Importantly, the appearance of carboxylated fIX_-18/41_ versus fIX in the V255M complex was very different than wild type complex: the fIX_-18/41_ was carboxylated by the V255M complex at earlier times and produced a ratio of fIX_-18/41_ to full length fIX of ∼4, which was 14-fold higher than the ratio for wild type complex (0.25, Fig. 5H). These results show that the processivity of the V255M carboxylase is impaired.

Carboxylation of fIX in the S300F complex was barely detectable (Fig. 5F). The extent of carboxylation was less than what was observed in the assay that assessed catalysis (Fig. 4B), which may reflect the different levels of KH_2_ in the two assays (150 µM in Fig. 4 to assay catalysis vs. 10 µM in Fig. 5 to assay processivity). S300F carboxylation of the fIX_-18/41_ challenge peptide was also poor.

### Processive MGP carboxylation is impaired in both V255M and S300F carboxylases

Processive carboxylation of MGP was also studied, using MGP_flag_-carboxylase complexes and MGP_1-64_ as the challenge peptide (Fig. 6A). The complexes were formed by co-infecting insect cells with baculoviruses containing MGP_flag_ and wild type or mutant carboxylases. The complexes were separated from free carboxylase by immunoadsorption onto anti-FLAG antibody resin. Complexes were incubated with a 50-fold excess of MGP_1-64_, followed by gel electrophoresis and PhosphorImager analysis. The results with the MGP_flag_-wild type carboxylase were very similar to those obtained with fIX: carboxylation of MGP_flag_ in the complex was much higher than MGP_1-64_ at early times in the reaction (Figs. 6B, E). Carboxylation of MGP_flag_ in the complex reached a plateau by the end of the reaction. This plateau represented full MGP_flag_ carboxylation, as determined by comparing the amount of Gla produced to the levels of the MGP_flag_-wild type carboxylase complex. The amount of complex was determined by performing an epoxidase assay to quantitate the carboxylase. Therefore, the wild type carboxylase processively modifies both MGP and fIX, even though these two VKD proteins have significant structural differences in the juxtapositioning of the EBD and Gla domains (e.g. Fig. 5A vs Fig. 6A).

**Figure 6.**
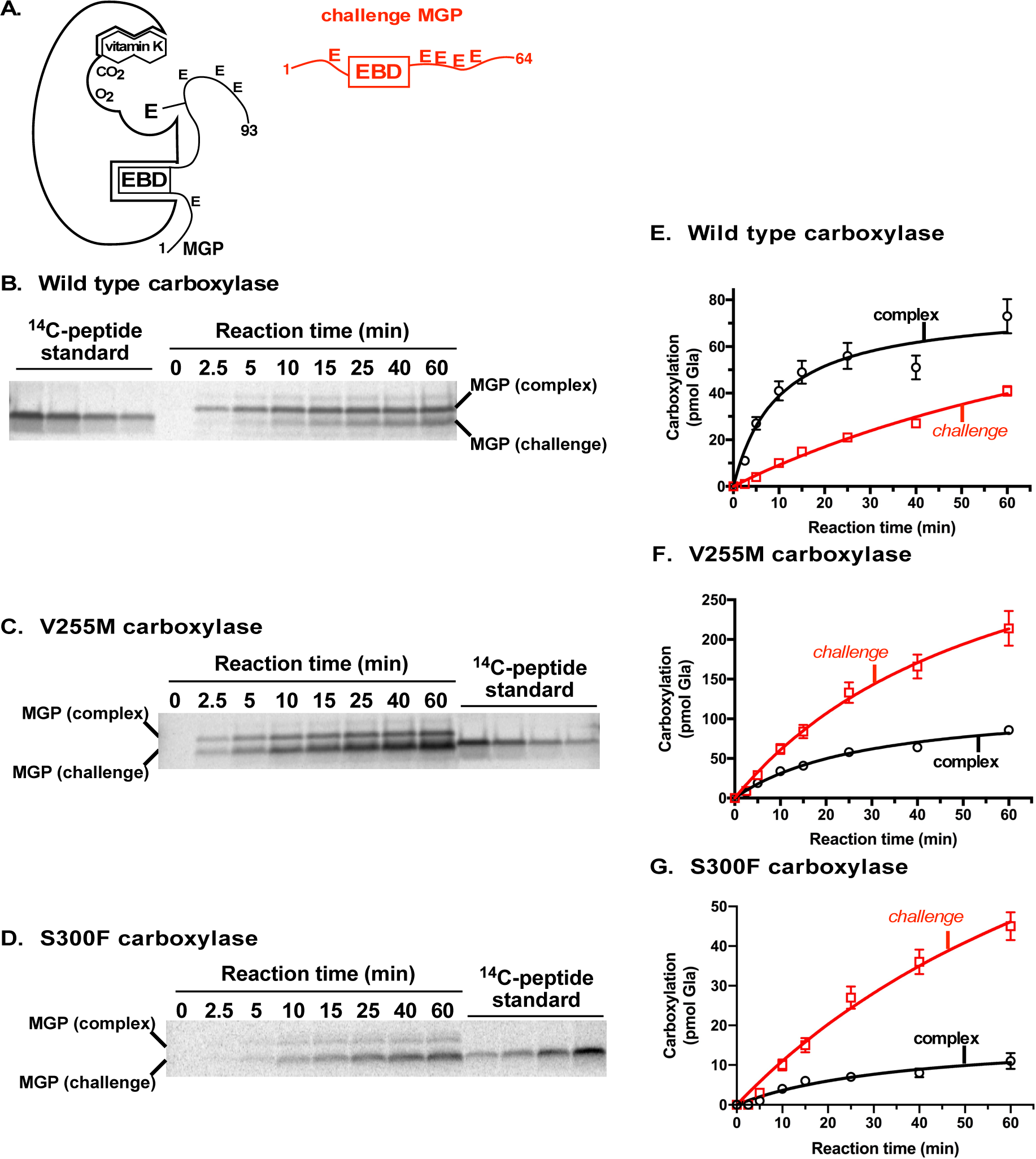
**A.** A reaction containing MGP_flag_-carboxylase complex and a 50-fold excess of the MGP_1-64_ challenge peptide was performed, and the carboxylation of the MGP_flag_ in the complex and MGP_1-64_ were monitored. The MGP_flag_-carboxylase complex was generated and purified as described in the legend to Figure 4. The MGP_1-64_ challenge peptide is the same peptide analyzed for carboxylation in Fig. 3C. **B-D.** The complexes between MGP_flag_ and wild type (B) or V255M (C) or S300F (D) carboxylase were incubated in reactions containing ^14^C-CO_2_, KH_2_ and MGP_1-64_, the products were analyzed by gel electrophoresis and PhosphorImager. **E-G.** Duplicate gels were quantitated to monitor MGP_flag_ and MGP_1-64_ carboxylation by wild type (E), V255M (F) and S300F (G) carboxylase variants.

Carboxylation was distinctly different for MGP_flag_-V255M versus MGP_flag_-wild type carboxylase complexes. The ratio of carboxylated challenge peptide to MGP_flag_ in the complex was much higher for V255M (Fig. 6C, F) than wild type carboxylase (Fig. 6B, E). As with the fIX experiments, the V255M complex produced a ratio of carboxylated challenge peptide to carboxylated full length protein of ∼4, which was 14-fold higher than the wild type complex. Therefore, the V255M carboxylase is also impaired for processive carboxylation of MGP.

Interestingly, despite the large amounts of carboxylated MGP_1-64_ generated by the V255M carboxylase, the MGP_flag_ in the complex still became fully carboxylated. Equal amounts of MGP_flag_-V255M and MGP_flag_-wild type carboxylase complexes were analyzed, and the amount of carboxylated MGP in the complex was the same (compare carboxylation of complex in Figs. 6E and 6F). An independent experiment was therefore performed to determine the amount of MGP_flag_-carboxylase complex that existed at the end of the reaction. This determination was possible because the complex had been immobilized on anti-FLAG resin. The challenge assay was performed as in Fig. 6, except that aliquots were also tested for the amount of carboxylase still bound to the anti-FLAG resin, and therefore to MGP_flag_. Most (∼80%) of the V255M and wild type carboxylases were still in a complex with MGP_flag_ at the end of the reaction (data not shown). Thus, while the challenge MGP_1-64_ was carboxylated, it did not displace the MGP in the complex.

The MGP_flag_-S300F carboxylase complex showed low levels of carboxylation of MGP_flag_ in the complex (Fig. 6D, G). However, the MGP_flag_-S300F carboxylase complex produced higher levels of MGP_1-64_ carboxylation, even at early times in the reaction. At the end of the reaction, the amount of carboxylated MGP_1-64_ challenge peptide was five times more than MGP_flag_ in the complex. The amount of carboxylated MGP_1-64_ produced by the S300F complex was less (20%) than that produced by the V255M complex (40 pmol versus 200 pmol, respectively), but nonetheless was a significant amount.

## DISCUSSION

How mutations in the gamma-glutamyl carboxylase cause the PXE-like disease was previously unknown, and these studies reveal the first mechanism for this defect, which is impaired processivity. S300F and V255M, which were previously identified in a patient with the PXE-like disease (15), were analyzed. The mutants were studied using large peptides that contain a Gla domain that undergoes multiple Glu to Gla conversions, and the EBD that tethers the VKD proteins to the carboxylase for processive carboxylation (10,13). Two substrates were tested: one derived from a hemostatic protein (fIX) and one derived the calcification regulator MGP. The V255M carboxylase variant showed higher levels of carboxylation than the wild type carboxylase with both substrates, while the S300F variant had low activity (Fig. 3). The results with the V255M variant suggested faster release of carboxylated products, which could impact processivity and lead to the release of incompletely modified protein. As described in the Introduction, we had previously shown that wild type carboxylase processivity results in full carboxylation of the Gla domain via EBD tethering of the VKD protein to the carboxylase (10). This property was revealed using a challenge assay in which the carboxylation of a fIX-carboxylase complex is monitored in the presence of an excess amount of a challenge protein (10). This approach was extended to studying the S300F and V255M mutant carboxylases. Both fIX and MGP were processively modified by wild type carboxylase (Figs. 5, 6), and the results indicate that tight complexes were formed that blocked the access of the challenge peptide and allowed full carboxylation of the VKD protein in the complex (Fig. 7A). In contrast, the complexes formed with the mutants were promiscuous, and allowed access of the challenge peptides into the active site (Fig. 7B). The V255M carboxylase produced a substantial amount of carboxylated challenge peptide in both the fIX- and MGP-containing complexes (Figs. 5H and 6F). The S300F carboxylase also showed impaired processivity in the MGP-S300F complex (Fig. 6D); however, the amount of product was ∼10% of that produced by the V255M carboxylase (Fig. 3C). Val255 and Ser300 are both in regions of the carboxylase with no assigned function (Fig. 1C), and the results indicate that these residues are important for maintaining a tight complex that ensures processivity.

**Figure 7.**
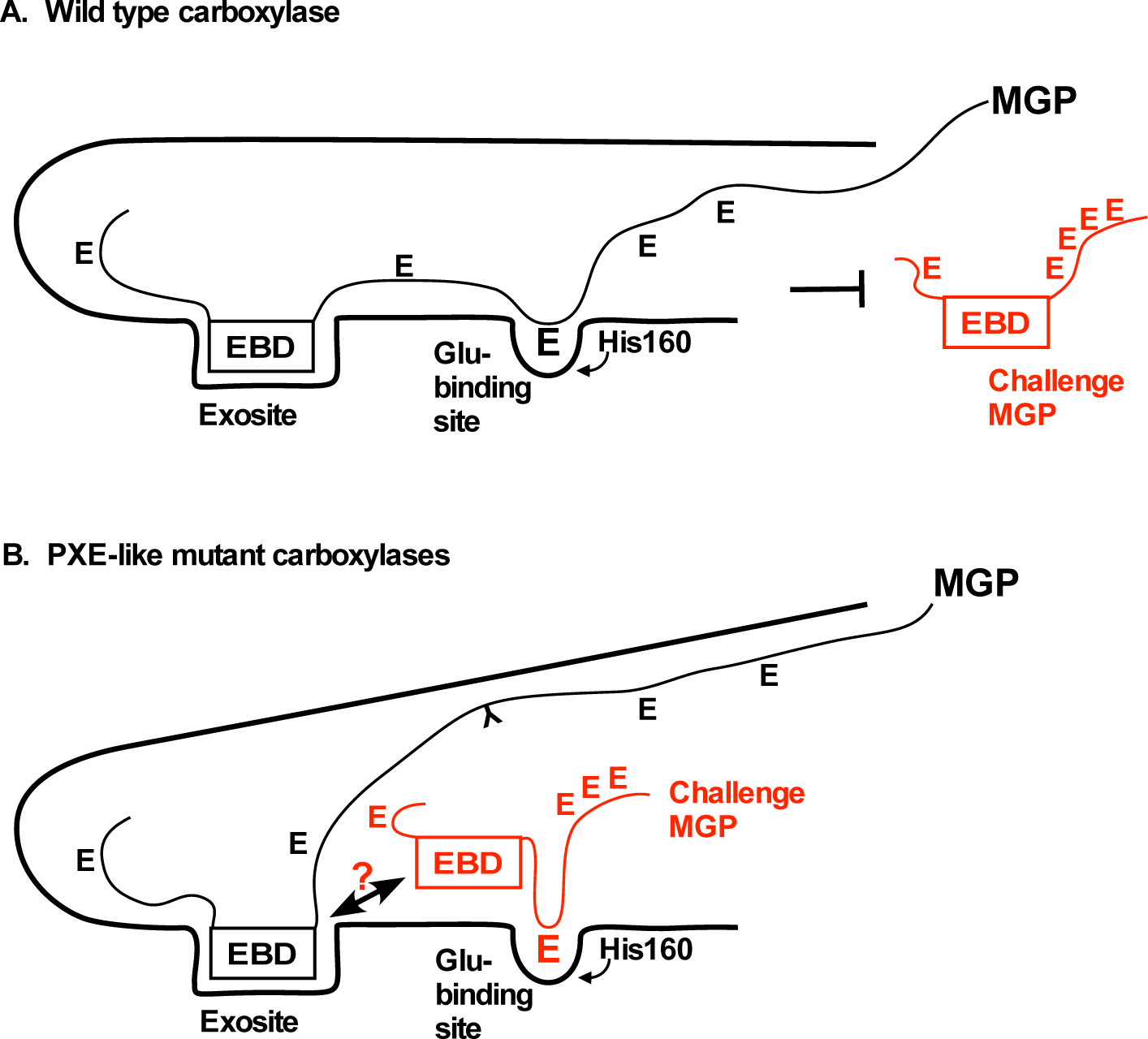
Impaired processivity in the PXE-like carboxylase variants V255M and S300F results in unregulated access of VKD proteins into the active site. **A.** Processive carboxylation involves tethering of the exosite binding domain (EBD) in VKD proteins to the carboxylase, which results in the carboxylation of multiple Glu residues to Gla’s in the Gla domain. The wild type carboxylase binds VKD protein substrates in a tight complex that blocks access of external VKD proteins into the active site. **B.** The V255M and S300F carboxylase variants bind VKD protein substrates in a promiscuous complex that allows access of other VKD proteins. Inappropriate access of the challenge VKD protein may not result in full carboxylation, generating a defective product. As described in the text, carboxylation is observed at low concentrations of the challenge VKD protein, which therefore requires high affinity binding to the carboxylase. High affinity binding is thought to occur through the exosite, which would displace the MGP in the complex and decrease its level of carboxylation. As decreased MGP carboxylation in the complex was not observed, these results raise the possibility of a second site of high affinity interaction between the carboxylase and VKD proteins.

Processivity was impaired with both fIX and MGP, raising the question as to why the patient has a mild bleeding defect and strong calcification defect. An open structure in the VKD protein-carboxylase complex (Fig. 7B) could generate a partially carboxylated product. Hemostatic factors that are partially carboxylated may retain activity, while partial modification of MGP may not be sufficient for function. Studies of fIX have shown that not all the Gla residues are required for activity (20). Similar studies have not been reported for MGP, which is thought to regulate calcification through the inhibition of bone morphogenic proteins (21). An alternative possibility is that incompletely carboxylated MGP may interfere with the regulation of calcification, while undercarboxylated clotting factors may have no impact on hemostasis. Future studies assessing why clotting and calcification are differentially impacted will be of interest in elucidating the physiological consequences of impaired carboxylase processivity.

The results from this work are important regarding the mild PXE-like phenotype observed in the proband’s mother and aunt. These relatives are heterozygous for the wild type and V255M carboxylase alleles. They are also heterozygous for wild type and p.R1141X ABCC6 alleles, which led to the proposal that partial loss of carboxylase and ABCC6 gene function resulted in a digenic disease producing mild PXE-like (15). However, the results with the V255M carboxylase variant suggest an alternative explanation. The V255M carboxylase had the surprising property of producing much more carboxylated challenge peptide than carboxylated MGP in the complex (Figs. 6C, F). Therefore, despite the presence of a wild type gene in the mother and aunt, these individuals may have a large amount of defective MGP produced by the V255M carboxylase, which may lead to the mild PXE-like disease. The proband shows increased severity of the disease, which may be due to a higher proportion of the MGP being defective because the proband lacks wild type carboxylase and instead has the S300F variant with impaired processivity and activity (Fig. 6G). The father and brother are heterozygous for the wild type and S300F carboxylase alleles, and for the wild type and p.R1141X ABCC6 alleles, but do not exhibit PXE-like symptoms (15). Multiple possibilities may account for the lack of disease, such as gender differences, or the reduced amount of product made from the S300F carboxylase variant (Fig. 3).

The results are relevant to whether vitamin K supplementation might help alleviate PXE-like disease symptoms. Vitamin K administration is valuable in a different disease caused by carboxylase mutations, i.e. vitamin K clotting factor deficiency 1 (VKCFD1)(22-24), where supplementation improves hemostasis. Vitamin K supplementation is thought to be effective because VKD protein binding to the carboxylase increases its affinity for vitamin K, and the carboxylase mutations may disrupt these interactions, resulting in the need for more vitamin K. The V255M mutant showed a response to vitamin K with fIX: the rate of V255M and wild type carboxylation was the same at high vitamin K concentrations (Fig. 4D), while V255M activity was 10% that of wild type carboxylase at low vitamin K levels (Fig. 5D). However, MGP carboxylation by V255M was the same as wild type carboxylase irrespective of vitamin K concentrations (Figs. 4E, 6E, 6F). These results suggest that vitamin K supplementation might not improve MGP carboxylation in the PXE-like disease. Other VKD proteins are also relevant to calcification, i.e. osteocalcin and Gla-rich proteins, which may respond differently than MGP to vitamin K levels.

The results from this work also shed light as to why patients on warfarin do not develop PXE-like symptoms. The warfarin patients have wild type carboxylase, and therefore a tight complex that should not allow inappropriate VKD protein access and the production of a defective product. However, warfarin treatment could cause the production of an undercarboxylated VKD protein by a different mechanism. Warfarin inhibits VKORC1, which provides reduced vitamin K to the carboxylase. VKORC1 is rate-limiting for carboxylation, and warfarin therefore decrease catalysis of the multiple Glu residues (25,26). While catalysis is normally much faster than release, the relative rates may change during warfarin therapy and result in premature release of partially carboxylated VKD protein. The type of undercarboxylated protein produced during warfarin therapy may not be the same as what is produced in PXE-like disease, which could explain why warfarin patients do not exhibit PXE-like symptoms.

The observation that the MGP substrates in the MGP-V255M carboxylase and MGP-wild type carboxylase complexes were both fully carboxylated (Figs. 6E, F) was surprising, considering what is known about the mechanism of carboxylation. The EBD is important in conferring high affinity binding of the carboxylase to Glu residues. When the EBD is provided as a free peptide, mM concentrations of Glu-containing peptide are required for carboxylation (as in Fig. 2). However, when the EBD and Glu residues are covalently linked, carboxylation occurs at µM concentrations (as in Fig. 3). In the challenge assays, the challenge peptides were efficiently carboxylated at µM levels (Figs. 5, 6). These observations suggest that the challenge peptide interacts with the carboxylase exosite; however, this interaction would displace the MGP in the complex and lower its level of carboxylation. Therefore, the observation that MGP in the V255M- and wild type carboxylase complexes was the same suggests that MGP in the complex remained bound to the carboxylase (Fig. 7B). The results suggest a second site for carboxylase-VKD protein interaction. A second site had previously been implicated by studies on osteocalcin, where the mature protein lacking the EBD has been shown to exhibit high affinity binding to the carboxylase (16). The possibility of a second site is also supported by the observation that mature clotting factors lacking the EBD bind to the carboxylase (27). Future studies that address how VKD proteins bind the carboxylase will be of interest, as VKD proteins with widely varying EBD affinities (12) must compete for a single carboxylase. As demonstrated by phenotypes of carboxylase mutations, how this diverse array of VKD proteins are appropriately modified in different tissues has important implications for human health and disease.

## EXPERIMENTAL PROCEDURES

### Preparation of purified carboxylase variants and complexes

Baculovirus containing wild type carboxylase, the V255M variant, or the S300F variant, in each case with a C-terminal alanine linker followed by a FLAG epitope (AAADYKDDDDK), was used to transfect SF21 cells (6 × 10^8^) with a multiplicity of infection of 5 as described (28,29). For preparation of wild type carboxylase-FLAG/factor IX complexes, SF21 cells (6 × 10^8^) were co-infected with baculoviruses using multiplicities of infection of 2 and 10, respectively, while for preparation of V255M-FLAG/factor IX or S300F-FLAG/factor IX complexes, the multiplicities of infection were 1 and 5, respectively. For preparation of untagged wild type, V255M, or S300F carboxylase complexes with N-terminally FLAG-tagged matrix Gla protein, SF21 cells (6 × 10^8^) were co-infected using a multiplicity of infection of 2 and 5, respectively.

For purified FLAG-tagged carboxylase variants, microsomes from infected SF21 cells were resuspended in 50 mM HEPES pH 7.4/500 mM NaCl/10% glycerol and adjusted to a total protein concentration of 4 mg/mL using a BCA assay (Pierce), then solubilized by addition of CHAPS (0.5%) and phosphatidyl choline (0.1%) followed by nutation at 4 °C for 1 h. After centrifugation (100000 × *g*, 4 °C, 1 h), aliquots of supernatant (1 ml) were nutated overnight with 100 μl anti-FLAG resin (Sigma, 0.6 mg/ml) at 4 °C. The resins were washed with five rounds of centrifugation (10000 × *g*, 1 min) and resuspension in 1 mL Buffer A (25 mM Tris pH 7.4/0.3% CHAPS/0.2% phosphatidyl choline/500 mM NaCl), all at 4 °C, followed by elution of carboxylase by nutation in 200 μl Buffer A with 100 μg/mL FLAG peptide (Sigma) for 1 h at 4 °C and centrifugation (10000 × *g*).

For FLAG-tagged carboxylase variants in complex with factor IX, microsomes from co-infected SF21 cells were resuspended in 50 mM HEPES pH 7.4/200 mM NaCl/10% glycerol, adjusted to a total protein concentration of 4 mg/ml, and solubilized as above. Aliquots (500 μl) were nutated overnight at 4 °C with 100 µl resin coupled to a monoclonal antibody (ESN1, 1 mg/ml, American Diagnostica) that recognizes uncarboxylated factor IX but not factor IX - 18/+41 (10). The resins were washed with five rounds of centrifugation (1000 × *g*) and resuspension in Buffer A, and resin-bound complexes were immediately used as described below.

For untagged carboxylase variants in complex with FLAG-tagged matrix Gla protein, microsomes from co-infected SF21 cells were applied to antiFLAG resin and washed as above for purified FLAG-tagged carboxylase variants. The washed resin-bound complexes were immediately used as described below.

### Preparation of [^14^C]fIX -18/+41 standards

Purified wild type carboxylase (∼100 pmols) was incubated in a cocktail (500 µL) containing 500 mM NaCl, 50 mM BES pH 6.94, 2.5 mM DTT, 0.16% CHAPS, 0.16% phosphatidyl choline, 200 µM KH_2_, and 1.3 mM [^14^CO_2_] (Perkin Elmer) for 6 hours, followed by addition of further KH_2_ (200 µM) and incubation for 12 hours. Ice-cold 20% trichloroacetic acid (500 µl) was added and the reaction was placed at 4 C for 24 h. The sample was centrifuged (10000 × *g*, -10 °C, 15 min) and the precipitate was twice washed with 500 µL acetone with incubation at -20 °C for 15 min, then the precipitate was resuspended in 25 mM ammonium bicarbonate pH 7.4/50% acetonitrile. The entire sample was injected on a Grace-Everest C18 column (150 × 2.1 mm, 5 µ) which underwent gradient elution from 25 mM ammonium bicarbonate pH 7.5/5 mM tetrabutylammonium chloride (solvent A) to 25 mM ammonium bicarbonate pH 7.5/5 mM tetrabutylammonium bicarbonate/50% acetonitrile (solvent B). The gradient conditions were as follows: flow rate 0.2 mL/min; 0-5 min: linear gradient from 2% solvent B to 70% solvent B; 5-30 min: linear gradient from 70% solvent B to 85% solvent B; 30-45 min: 85% solvent B. Separation of carboxylated fIX -18/+41 from uncarboxylated was effected by the tetrabutylammonium chloride ion pairing reagent and was monitored using absorbance at 210 nm. Fractions containing [^14^C]fIX -18/+41 were pooled, lyophilized to a small volume, and analyzed by scintillation counting to determine counts per unit volume.

### Assessment of activity of carboxylase variants with varied activation and substrate peptides

Purified carboxylase (∼25 pmols) was incubated in a cocktail (∼160 μl) containing 500 mM NaCl, 50 mM BES pH 6.94, 2.5 mM DTT, 0.16% CHAPS, 0.16% phosphatidyl choline, 200 μM KH_2_, and 1.3 mM [^14^C]CO_2_ for various times at 25 °C. When present, the factor X exosite binding domain (EBD) peptide (SLFIRREQANNILARVTR) was present at 10 μM or, in one experiment, at a range of concentrations, while in one experiment the matrix Gla protein EBD peptide (NPFINRRNANTFISPQQR) was present at a range of concentrations. Substrate peptides, when present, were as follows: the pentapeptide FLEEL (Anaspec), 2.5 mM; fIX -18/+41 (the propeptide and Gla domain of factor IX, Anaspec), 1 uM; MGP 1-64 (the first 64 amino acids of matrix Gla protein, Anaspec), 1 uM. FLEEL carboxylation was assessed by measurement of [^14^C]CO_2_ incorporation into substrate peptides as described (29).

For assessment of factor IX -18/+41 carboxylation, reaction aliquots underwent chloroform/methanol precipitation (30) of the substrate peptide. Dried precipitates were resuspended in 100 μl 25 mM ammonium bicarbonate/2 mM DTT and the resuspended samples were analyzed by HPLC with a C18 column (Grace-Everest, 150 × 2.1 mm, 5 μ particle size) with gradient elution and analysis as above. Fractions containing carboxylated factor IX - 18/+41 were pooled and [^14^C]CO_2_ incorporation into the substrate peptide was determined by scintillation counting.

For assessment of MGP 1-64 carboxylation, reaction aliquots were quenched by addition to SDS/PAGE loading dye and samples were loaded onto a Criterion 16.5% TRIS/TRICINE gel (BioRad) along with [^14^C]BSA counting standards (American Radiolabelled Chemicals, further purified by passage over a P100 size exclusion column) and [^14^C]-labelled molecular weight markers (Perkin Elmer). Gels were electrophoresed, then proteins and peptides were transferred to nitrocellulose membranes (0.2 micron, GE), and [^14^C] counts were imaged and quantized by phosphoimaging of the nitrocellulose membranes as described (10) followed by comparison of radioactivity incorporated into factor IX -18/+41 to the BSA standards.

### Single turnover assay for complexed factor IX carboxylation

Factor IX/carboxylase complexes bound to ESN1 resin (100 μl) were resuspended in a reaction cocktail (350 μl) consisting of 500 mM NaCl, 50 mM BES pH 6.94, 2.5 mM DTT, 0.16% CHAPS, 0.16% phosphatidyl choline, 1.3 mM [^14^C]CO_2_, and 200 μM KH_2_ and the reactions were nutated at 24 °C. At various time points, aliquots were withdrawn and quenched by addition to SDS/PAGE gel dye followed by boiling (5 min) and centrifugation (700 × *g*) to pellet resin. Aliquots were analyzed by SDS/PAGE using an 8% TRIS gel, in each case along with [^14^C]BSA standards and [^14^C]-labelled molecular weight markers. Proteins were then transferred to 0.45 micron nitrocellulose membranes (GE) and [^14^C] counts were imaged and quantized as above. Resin-bound factor IX/carboxylase complex was also assessed in parallel for vitamin K epoxidation activity using a similar cocktail but including FLEEL (2.5 mM) and factor X EBD (10 μM). Epoxidation activity was measured by addition of 10 µl K25 standard (GL Synthesis, 2 nmol) followed by extraction of reaction aliquots with 3 volumes 1:1 ethanol:hexane. The organic phase was evaporated to dryness and the extracts were resuspended in 60 µl ethanol and analyzed for vitamin K epoxide using a HPLC assay as described (31).

### Single turnover assay for complexed matrix Gla protein

Matrix Gla protein/carboxylase complexes bound to antiFLAG resin (100 μl) were resuspended in a reaction cocktail (250 μl) containing 500 mM NaCl, 50 mM BES pH 6.94, 2.5 mM DTT, 0.16% CHAPS, 0.16% phosphatidyl choline, 1.3 mM [^14^C]CO_2_, and 200 μM KH_2_ and the reactions were nutated at 24 °C. Carboxylation of matrix Gla protein was determined as for the factor IX single turnover assay except for use of a Criterion 16.5% TRIS/TRICINE gel (BioRad) with [^14^C]fIX -18/+41 standards for quantization, and transfer to 0.2 micron nitrocellulose membranes (GE). Vitamin K epoxidation activity of resin-bound complexes was determined in parallel as above.

### Challenge assays for complexed factor IX carboxylation

Factor IX/carboxylase complexes bound to ESN1 resin (100 μl) were resuspended in a reaction cocktail (350 μl) consisting of 500 mM NaCl, 50 mM BES pH 6.94, 2.5 mM DTT, 0.16% CHAPS, 0.16% phosphatidyl choline, 1.3 mM [^14^C]CO_2_, 1 μM factor IX -18/+41, and 10 μM KH_2_ and the reactions were nutated at 24 °C. At various time points, two aliquots were withdrawn and quenched by addition to SDS/PAGE gel dye followed by boiling (5 min) and centrifugation (700 × *g*) to pellet resin. One aliquot from each time point was analyzed by SDS/PAGE using a Criterion 16.5% TRIS/TRICINE gel (BioRad) along with [^14^C]fIX -18/+41 standards, while the second aliquot was analyzed by SDS/PAGE using an 8% TRIS gel along with [^14^C]BSA standards, in each case with [^14^C]-labelled molecular weight markers. Peptides and proteins were then transferred to 0.2 micron nitrocellulose membranes (GE) and [^14^C] counts were imaged and quantized as above. Resin-bound factor IX/carboxylase complex was also assessed in parallel for vitamin K epoxidation activity using a similar cocktail but with 200 μM KH_2_ and with factor IX -18/+41 replaced by FLEEL (2.5 mM) and factor X EBD (10 μM) followed by organic extraction and HPLC assay as above.

A further challenge assay was also performed exactly as above except that fIX -18/+41 was replaced by an equivalent concentration of MGP 1-64.

### Challenge assay for complexed matrix Gla protein carboxylation in the presence of MGP 1-64

Matrix Gla protein/carboxylase complexes bound to antiFLAG resin (100 μl) were resuspended in a reaction cocktail (250 μl) containing 500 mM NaCl, 50 mM BES pH 6.94, 2.5 mM DTT, 0.16% CHAPS, 0.16% phosphatidyl choline, 1.3 mM [^14^C]CO_2_, 1 μM MGP 1-64, and 10 μM KH_2_ and the reactions were nutated at 24 °C. Carboxylation of matrix Gla protein and MGP 1-64 was determined as for the factor IX challenge assay except for use of a single aliquot at each time point that was analyzed using a Criterion 16.5% TRIS/TRICINE gel (BioRad). Vitamin K epoxidation activity of resin-bound complexes was determined in parallel as above.

A further challenge assay was also performed exactly as above except that MGP 1-64 was replaced by an equivalent concentration of fIX -18/+41.

### Western analysis of carboxylase variants, factor IX, and matrix Gla protein

Purified carboxylase variants underwent SDS/PAGE alongside BAP-FLAG quantization standards (Sigma) using a Criterion 10% TRIS gel (BioRad) followed by transfer to a 0.45 micron nitrocellulose membrane (GVS North America). Membranes were analyzed using an antiFLAG antibody (Sigma, 0.4 μg/ml) followed by a goat anti-rabbit antibody conjugated to IR dye 800 CW (0.2 μg/mL, LiCor Biosciences). Carboxylase variants were quantized using an Odyssey LiCor scanner according to the manufacturer’s instructions.

Carboxylase (with or without FLAG tag) in microsome suspensions was analyzed as above except using a Novex BIS-TRIS 4-12% gel (Invitrogen) and an antibody against the C-terminus of the carboxylase (Pudota 1998) (0.4 µg/mL). Factor IX in microsome suspensions was also analyzed using Novex gels and a polyclonal antibody (0.4 µg/mL).

Matrix Gla protein microsome suspensions were also analyzed using Novex gels followed by transfer to a 0.2 micron nitrocellulose membrane (BioRad). Membranes were analyzed using an antiFLAG antibody (Sigma, 0.4 μg/ml) and secondary antibody as above, however all manipulations were performed using an altered buffer system composed of 10 mM TRIS pH 8/150 mM NaCl/0.05% TWEEN due to loss of signal with the typical buffer system.

## ACKNOWLEDGEMENTS

This work was supported by NIH grants HL055666 and HL081093 to KLB. KWR was supported by NIH grant AG051601 and NSF award 1516220.

## AUTHOR CONTRIBUTIONS

MAR, KWR and KLB designed the experiments. MAR and KWH performed all of the experiments. MAR, KWR, and KLB wrote the manuscript. KLB and KWR obtained funding for the research.

